# Differential expansion and retention patterns of LRR-RLK genes across plant evolution

**DOI:** 10.1101/2023.07.26.549740

**Authors:** Zachary Kileeg, Aparna Haldar, Hasna Khan, Arooj Qamar, G. Adam Mott

**Affiliations:** Department of Biological Sciences, University of Toronto Scarborough, 1265 Military Trail, Toronto, ON M1C 1A4, Canada; Department of Cell and Systems Biology, University of Toronto, 25 Willcocks Street, Toronto, ON M5S 3B2, Canada; Centre for the Analysis of Genome Evolution & Function, University of Toronto, 25 Willcocks Street, Toronto, ON M5S 3B2, Canada

## Abstract

To maximize overall fitness, plants must accurately respond to a host of growth, developmental, and environmental signals throughout their life. Many of these internal and external signals are perceived by the leucine-rich repeat receptor-like kinases, which play roles in regulating growth, development, and immunity. This largest family of receptor kinases in plants can be divided into subfamilies based on conservation of the kinase domain, which demonstrates that shared evolutionary history often indicates shared molecular function. Here we investigate the evolutionary history of this family across the evolution of 112 plant species. We identify lineage-specific expansions of the malectin-domain containing subfamily LRR subfamily I primarily in the Brassicales and bryophytes. Most other plant lineages instead show a large expansion in LRR subfamily XII, which in Arabidopsis is known to contain key receptors in pathogen perception. This striking asymmetric expansion may reveal a dichotomy in the evolutionary history and adaptation strategies employed by plants. A greater understanding of the evolutionary pressures and adaptation strategies acting on members of this receptor family offers a way to improve functional predictions for orphan receptors and simplify identification of novel stress related receptors.

## Introduction

In order to survive, plants must appropriately respond to numerous extracellular signals. These signals guide both development and immunity in the plant. To aid in discriminating between these signals, plants have evolved a large suite of cell surface receptors. Many of these receptors are receptor-like kinases (RLKs), with the largest group in Arabidopsis (*Arabidopsis thaliana*) being the leucine-rich repeat receptor-like kinase family (LRR-RLK) (Shiu and Bleecker, 2001; Lehti-Shiu et al., 2009).

The LRR-RLKs have three protein domains: an N-terminal extracellular domain (ECD) to receive signals and mediate the physical interactions that lead to the formation of receptor complexes, a single transmembrane domain (TM), and a C-terminal intracellular kinase domain (KD). Previous analyses have divided the LRR-RLKs into numerous subfamilies based on an inferred phylogeny from alignment of their KDs(Liu et al., 2017; Shiu and Bleecker, 2001). The LRR-RLKs are named for the presence of numerous leucine-rich repeats (LRR) within the ECD. In addition to these canonical repeats, the ECDs of LRR-RLK-I (LRR-I), LRR-RLK-VIII.1 (LRR-VIII.1) and LRR-RLK-VIII.2 (LRR-VIII.2) subfamily members contain malectin, or malectin-like, domains (MD/MLDs). MDs show sequence similarity to the malectins, which are small ER-resident disaccharide-binding proteins in animal cells (Schallus et al., 2008, 2010), while the MLD is a plant specific domain composed of two tandem MDs.

As they often impact agronomically important traits, the LRR-RLKs have long been seen as attractive targets for crop improvement (Diévart and Clark, 2004; Lemmon et al., 2018; Rodríguez-Leal et al., 2017). In spite of this interest, only a minority of LRR-RLKs have defined biological functions. The majority of LRR-RLKs that have been characterized to date are broadly involved in either immunity or growth and development (Tang et al., 2010). An impediment to functional characterization is the large number of closely related genes with the potential for high rates of functional redundancy (Rodriguez-Leal et al., 2019; Nimchuk et al., 2015).

The expansion and diversification of LRR-RLK repertoires within plant lineages has resulted from both whole genome duplications and smaller segmental or tandem duplications (Panchy et al., 2016). Whole genome duplication has occurred many times over the course of angiosperm evolution (Soltis et al., 2009; Renny-Byfield and Wendel, 2014), while tandem and segmental duplications are also common (Rizzon et al., 2006). Interestingly, rates of LRR-RLK duplication and retention appear correlated to their biological function (Lehti-Shiu et al., 2009). Specifically, subfamilies involved in stress responses are greatly expanded by tandem duplication, while developmental LRR-RLKs lack this signature (Shiu et al., 2004; Tang et al., 2010). The genes from expanded subfamilies also show evidence of increased positive selection when compared to LRR-RLKs from nonexpanded subfamilies (Tang et al., 2010). This selection is observed predominantly in the ECD, which is responsible for signal recognition (Tang et al., 2010).

In this work, we examine the expansion of the LRR-RLK family across plant evolution. We show a specific expansion of the LRR-I subfamily within the Brassicales that is driven mainly by tandem duplication. Most other plant lineages show expansion of the LRR-RLK-XII (LRR-XII) subfamily, which contains the best studied LRR-RLKs involved in pathogen perception and response. Interestingly, almost all plant species show a preferential expansion of either the LRR-I or LRR-XII family. We then further examine the diversity of LRR-I protein sequences, demonstrating two distinct domain architectures and the presence of a well-conserved cleavage motif across plant lineages.

## Methods

### Plant genomes and LRR-RLK identification

Representative protein sequences were retrieved from Phytozome version 13 (Goodstein et al., 2012) for the 112 plant and algae species available as of May 2020. These 112 species were assembled into an approximate species tree using the NCBI taxonomy browser (Schoch et al., 2020) with red algae (*Porphyra umbilicalis*, species code: Pumbi) as the outgroup. Each species was assigned a five-letter code for ease of identification (gene codes and species names can be found in Supplementary Table 1) and divided into taxonomic groups for downstream analysis. Species were assigned to their respective orders within angiosperms where appropriate, with mosses and liverworts assigned to the bryophytes, and green algae to Chlorophyta. Orders containing two or fewer species were grouped together into a single group termed ‘Other’.

To identify LRR-RLK genes from the list of representative protein sequences we first identified proteins that contained an LRR domain using predict-phytoLRR (Chen, 2021), a protein kinase domain (PF00069, PF07714), or an NB-ARC (PF00931) domain using hmmsearch (Eddy, 2011) (v3.3). We included all proteins containing both an LRR and kinase domain, but lacking an NB-ARC domain in the collection of LRR-RLKs. The kinase domains of all LRR-RLK sequences were aligned using MAFFT (Katoh and Standley, 2013) (v.7.453) and an unrooted phylogeny was inferred using fasttree (Price et al., 2010) (v2.1.11) with default settings. Using this tree we assigned the LRR-RLKs to subfamilies using Arabidopsis LRR-RLK annotations as a guide (Wang et al., 2019). In cases where existing Arabidopsis subfamilies resolved into two or more clades, the smaller group of the subfamily was split and designated into a new division of that subfamily. We then calculated the proportion of LRR-RLKs belonging to each subfamily in each species.

### Analysis of tandem duplication

We identified any LRR-RLKs that arose via tandem duplication by manual inspection. We defined tandem duplicates as LRR-RLKs belonging to the same subfamily on the same chromosome separated by no more than two intervening genes with at least 50% similarity between their entire protein coding sequences.

### LRR-I and LRR-XII phylogenies

We inferred separate phylogenetic trees for LRR-I and LRR-XII subfamilies using sequences from all 112 species. The kinase domains of the LRR-RLK protein sequences were aligned using default settings in MAFFT (Katoh and Standley, 2013) (v.7.453) and maximum-likelihood phylogenies were constructed using the JTT+F+R10 substitution model based on modelfinder (Kalyaanamoorthy et al., 2017) in IQtree (Nguyen et al., 2015) with 1000 bootstrap replicates (v1.6.12). LRR-I trees were rooted using a variety of human and Arabidopsis kinase sequences (Sakamoto et al., 2012). Subclades were assigned based on a bottom-up approach where sequences from new branches were merged with the growing subclade if they shared an average of 60% sequence similarity and a bootstrap support of at least 90%. This was done until the next branch to be added contained sequence similarity <60%.

### Ancestral state reconstruction

We used ancestral state reconstruction to infer the maximum-likelihood number of LRR-I or LRR-XII subfamily members at each branch of the NCBI species tree using fastAnc from the R package phytools(Revell, 2012) (v0.7-70). The branch-specific expansion rate for each subfamily was calculated by dividing the number of LRR-RLKs at a given node by the number of LRR-RLKs of the same subfamily in the most recent ancestral branch.

### Motif analysis

We input the LRR-I protein sequences from all 112 species into memesuite (Bailey et al., 2015) (v5.4.1) and considered the top ten most significant results that contained at least 5 amino acids in the motif. The protein sequences were compared against the UniprotKB protein database to examine motif prevalence using ScanProsite (de Castro et al., 2006) (v20.0). Occurrences of exact motif matches were separated by subfamily and counted.

### Analysis of domain architecture

LRR-I protein sequences for each subclade were analyzed with HMMScan in HMMER (Eddy, 2011) (v.3.3) using the pfam-a HMM set (Mistry et al., 2021). All significant domain hits (E-value <1) were output and counted. Transmembrane regions were predicted using TMHMM (Krogh et al., 2001) (v2.0) and signal peptides were predicted using SignalP (Almagro Armenteros et al., 2019) (v5.0) for representative sequences. All predicted domains were compared and consensus domains were plotted using a representative sequence. Disulfide bonds were inferred from the predicted protein structure of the Arabidopsis FRK1 protein (AF-O64483-F1). Unique domains were plotted on the most common architecture in the general area that the domain appears.

### Protein structure mapping

The predicted protein structure of FRK1 (AF-O64483-F1) was obtained from AlphaFold (Jumper et al., 2021) (v2.0) and the transmembrane region and cytosolic region removed from the model in Pymol (The PyMOL Molecular Graphics System, Version 2.0 Schrödinger, LLC). The locations of motifs and other sequence features from MEME or HMMScan were mapped to the extracellular region in Pymol.

### Tissue Expression Analysis

Tissue expression analysis for 42 Arabidopsis LRR-I genes was performed using e-Northern (Toufighi et al., 2005) using default settings.

## Results

### LRR-RLKs display subfamily-specific expansion patterns across 112 plant and algal species

Based on their known domain structure, we identified LRR-RLK genes as those that encode for proteins with an LRR domain and a kinase domain, but lacking an NB-ARC domain. This definition was used to identify all putative LRR-RLK genes in 112 plant and algal genomes. The protein sequences of the kinase domains from all putative LRR-RLK genes for each species were aligned and subfamily designations were determined based on similarity to known Arabidopsis genes. The number of LRR-RLK genes present in each subfamily for each species was used to identify subfamily-specific rates of gene expansion and retention. We looked at the proportion of genes in each subfamily and found that the majority of subfamilies have consistent proportions across all plant species studied (Figure 1A). Six subfamilies deviate from the others, showing considerable variation between plant species and containing large numbers of expanded genes (Figure 1A, subfamilies LRR-I, LRR-III, LRR-VIII-1, LRR-VIII-2, LRR-XI, and LRR-XII). These subfamilies have the highest proportion of LRR-RLKs and display the greatest variability in gene proportion across the sampled species, bot expected characteristics of genes involved in stress responses.

**Figure 1.**
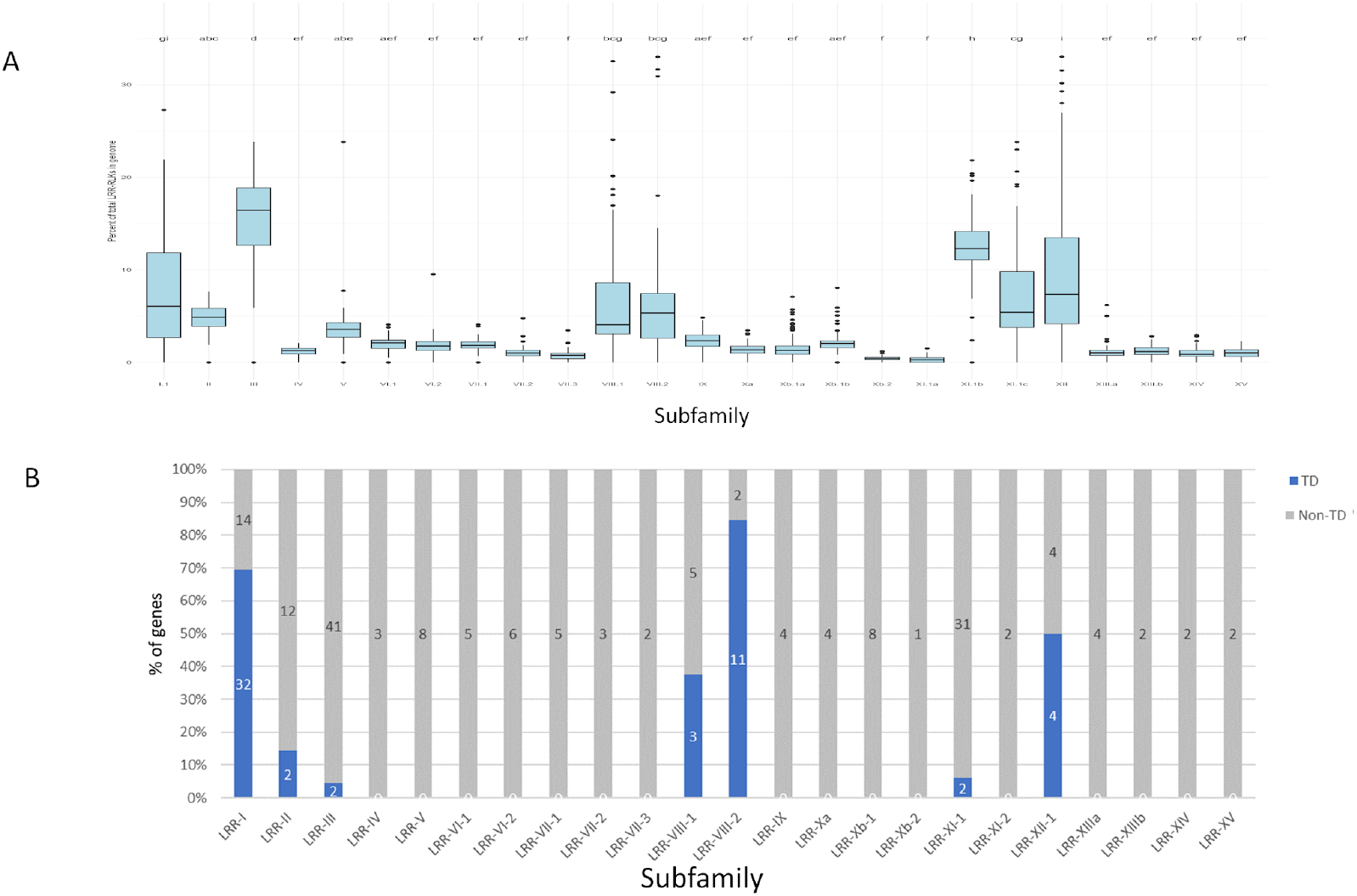
Expansion, duplication, and counts of LRR-RLKs across 112 plant and algae species. The LRR-RLK subfamily members have undergone asymmetric expansion (A). Subfamily expansion is represented as the percentage of LRR-RLKs belonging to each subfamily out of the total number of LRR-RLKs in each genome. Letters represent the significance group of each subfamily, as determined by Tukey’s HSD test at alpha = 0.05. Subfamilies with the shared letters are statistically similar to one another, while subfamilies with different letters are significantly different from one another. Species with less than 10 total LRR-RKs were omitted. Subfamilies LRR-I, LRR-III, LRR-VIII, and LRR-XI have a higher proportion of total LRR-RLKs compared to the other subfamilies, demonstrating increased diversification. LRR-RLKs have undergone tandem duplication (B). Blue bars represent tandem duplicated genes, while grey bars represent non-tandem duplicated genes. Tandem duplicates are those in the same subclade separated on a chromosome by no more than two intervening genes.

We next sought to determine the mechanism of subfamily expansion, as tandem duplication and subsequent retention is also correlated with stress response genes (Hanada et al., 2008; Qiao et al., 2019). In Arabidopsis, the expansions observed in subfamilies LRR-I, LRR-VIII-1, LRR-VIII-2, and LRR-XII are associated with high rates of tandem duplication (>50% of subfamily genes), suggestive of potential roles in stress responses (Figure 1B).

### The expansions of subfamilies LRR-I and LRR-XII are largely negatively correlated

Having identified several subfamilies with evidence of expansion through tandem duplication, we chose to focus on subfamily LRR-I. LRR-I was chosen because it is the largest subfamily within Arabidopsis, has extensive tandem duplication, and has members known to function in biotic stress responses (Yeh et al., 2016; Yuan et al., 2018; Asai et al., 2002). As a comparator, we chose subfamily LRR-XII, as it has a similar expansion pattern to LRR-I, shows high tandem duplication, and is known to contain receptors involved in the biotic stress response, including FLS2 (Zipfel et al., 2006; Chinchilla et al., 2006; Mott et al., 2016). To explore the lineage-based expansion dynamic of these subfamilies, we grouped the sequences from LRR-I and LRR-XII into their respective orders and inferred unrooted maximum-likelihood phylogenies of both (Supplemental Figure 1). This revealed an asymmetric expansion pattern, where Brassicales has a clear and particularly broad expansion in LRR-I but not in LRR-XII, whereas Poales shows the opposite trend (Supplemental Figure 1). To avoid any sampling effects arising from input species number, we explored this asymmetry by comparing the proportions of LRR-I and LRR-XII receptors across orders (Figure 2A). As expected, Poales has significantly more LRR-XII genes than LRR-I (Figure 2A).

**Figure 2.**
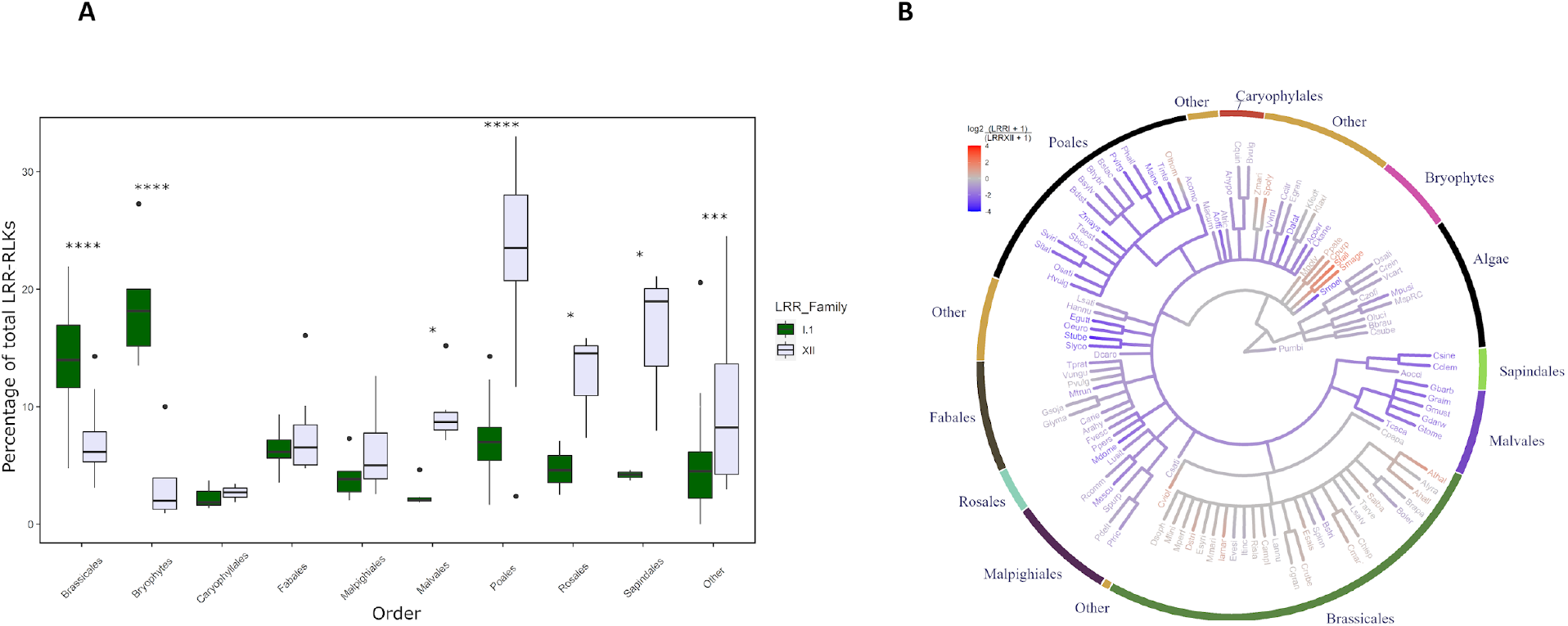
Comparison of LRR-I and LRR-XII receptor counts across different plant orders. The percentage of total LRR-RLKs found in subfamilies LRR-I and LRR-XII in each given order (A). Outliers (> Q3 + 1.5xIQR) are plotted as dots. Significance values represent an ANOVA followed by Tukey’s HSD. * = p<0.05, ** = p<0.01, *** = p<0.001, **** = p<0.0001. LRR-RLK subfamilies have different ratios of expansion across evolutionary time (B). Colours indicate the log of the ratio of LRR-I to LRR-XII genes at each node. Numbers at terminal nodes represent the ratio for those species and internal nodes represent the ratio for the common ancestor at that branch. Red algae (*P. umbilicalis)* is used as the species outgroup.

Similarly, the Malvales, Rosales, and Sapindales orders all have significantly more LRR-XII receptors than LRR-I (Figure 2A). In contrast, the opposite trend is seen in Brassicales and bryophytes, in which we identify significantly more LRR-I than LRR-XII (Figure 2A). The order Brassicales and the bryophytes are notable for their relative dearth of subfamily LRR-XII members compared to other plant lineages, suggesting a possible negative correlation between expansions in these two subfamilies.

We next tested whether this observed expansion asymmetry was broadly true in plants by comparing the ratio of LRR-I and LRR-XII gene count in each plant species (Figure 2B – external nodes). The majority of species show a large expansion in subfamily LRR-XII without a similar expansion in LRR-I (shown in blue). Fewer independent plant groups show the inverse pattern, which is like that observed in Arabidopsis and the bryophytes (red). We observe fewer lineages with similar expansions in these two subfamilies (grey). To estimate at what point during plant evolution these patterns emerged, we used ancestral state reconstruction to infer branch-specific expansion for each species (Figure 2B – internal nodes, Supplemental Figure 2). We found that the high number of LRR-I genes present in modern-day Brassicales and especially in the Brassicaceae family may be the result of two expansion events. The first is in the ancestor of Brassicales and a second in the ancestor of Brassicaceae, followed by local duplication events (Figure 2B, Supplemental Figure 2). Concomitant with this expansion of LRR-I, there is a reduction in the number of LRR-XII genes in the Brassicales. In contrast, numerous non-Brassicales angiosperms show expansion in LRR-XII rather than LRR-I (Figure 2). In particular, the order Poales shows the largest expansion of subfamily LRR-XII among any of the plant groups studied (Figure 2B, Supplemental Figure 2).

### There are twenty-five well-supported subclades within the LRR-I subfamily

Based on the preferential expansion of LRR-I in Brassicales and bryophytes, we chose to investigate the evolutionary relationship of LRR-I sequences. Of the 112 species of plant and algae we investigated, 101 contain at least one LRR-I gene. We used the sequences from these 101 species to reconstruct the phylogeny of LRR-I. We identified 25 subclades within LRR-I supported by high sequence similarity and widespread bootstrap support (Figure 3). Subclades LRR-I.1 through LRR-I.4 exclusively contain bryophyte sequences, while the remaining bryophyte sequences do not cluster in any well-supported subclades. Subclades LRR-I.5 through LRR-I.7, LRR-I.16, LRR-I.18, LRR-I.19, and LRR-I.24 contain sequences from a diverse range of plant species, but do not contain any bryophyte sequences (Figure 3). Notably, subclades LRR-I.1 through LRR-I.4, LRR-I.9 through LRR-I.15, LRR-I.17, LRR-I.20 through LRR-I.23, and LRR-I.25 are lineage-specific, with >95% of the sequences derived from a single order of plants (Figure 2). Among these, Brassicales and Poales have the largest and second-largest expansions, respectively (Figure 2). There also appears to be species-specific expansion, denoted by subclades LRR-I.1 and LRR-I.13 through LRR-I.15 although this may be a consequence of limited lineage sampling (Figure 3).

**Figure 3.**
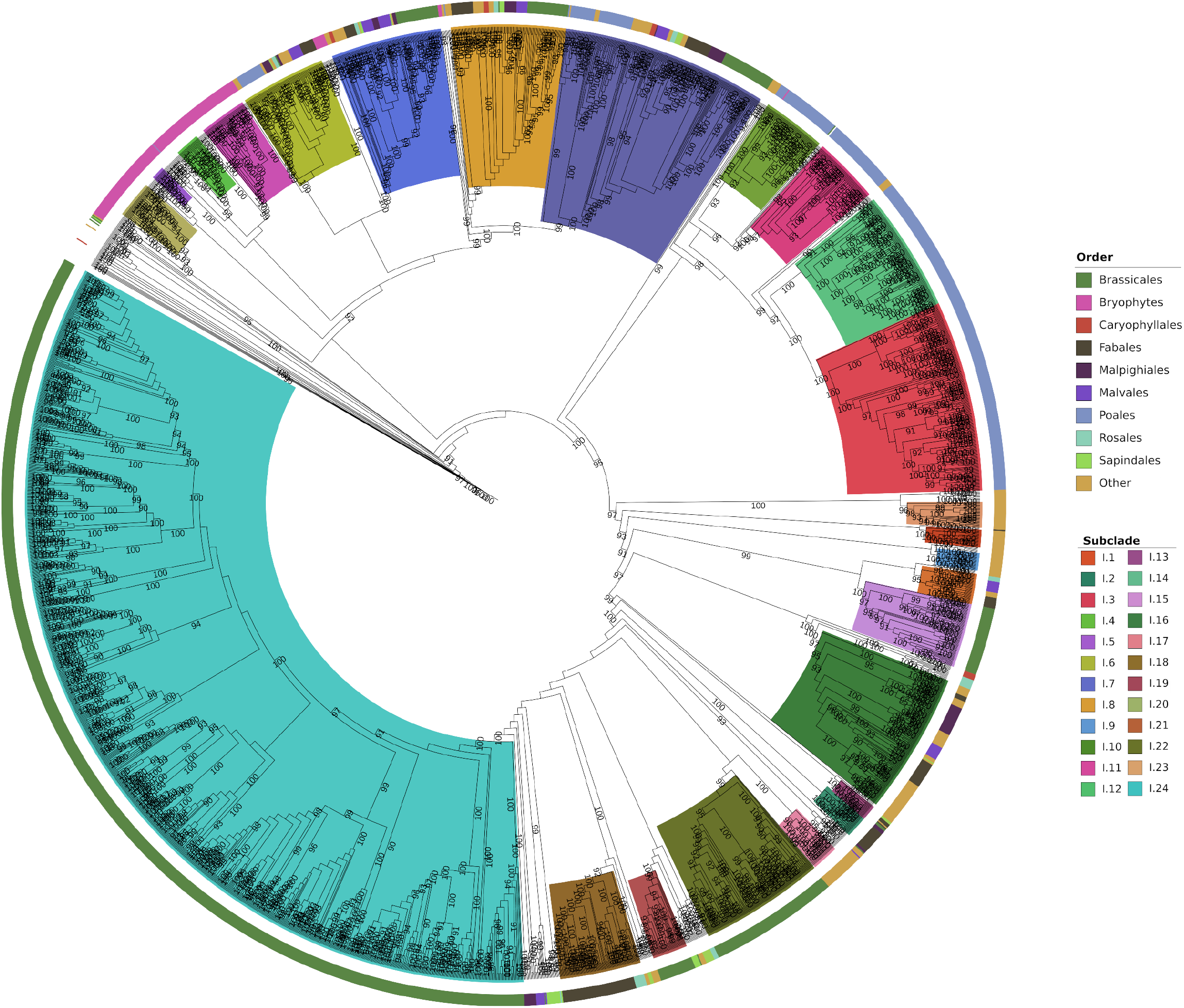
Maximum-likelihood phylogeny of LRR-I inferred from 112 species of plants and algae. Outer ring denotes the order each sequence falls into. Inner ring colours denote the subclade to which each sequence belongs. Bootstrap values represent 1000 re-samplings. Subclades were considered as such if sequences shared at least 60% sequence similarity to one another and the subclade contained bootstrap support >90%. Phylogeny was rooted using a variety of Arabidopsis and Human kinases.

### Novel motifs and domain architectures

After observing high levels of species and lineage-specific expansion and retention of LRR-I receptor repertoires, we hypothesized this may be evidence of diversification and neofunctionalization within this subfamily. We determined that LRR-I receptors primarily display one of two architectures: the first has an ECD containing a malectin-like domain and a variable number of leucine-rich repeats (2-4) followed by a transmembrane domain and a cytosolic kinase domain, while the second type is either missing the MLD and has a longer LRR domain or contains a truncated MLD (Figure 4A, Supplementary Figure 3C). The second architecture is found almost exclusively in the bryophyte-containing subclade LRR-I.1 (Figure 3). The angiosperm LRR-I proteins almost universally contain an MLD domain and display the first architecture, indicating a clear division between the bryophytes and flowering plants (Figure 4). Uncommon domains also were found in several regions including the N-terminal region of the MLD, between the repeated malectin domains within the MLD, within gaps between and overlapping with LRR domains, spanning the transmembrane/juxtamembrane region, and within the C-terminal region of the protein (Figure 4B, Supplemental Figure 3).

**Figure 4.**
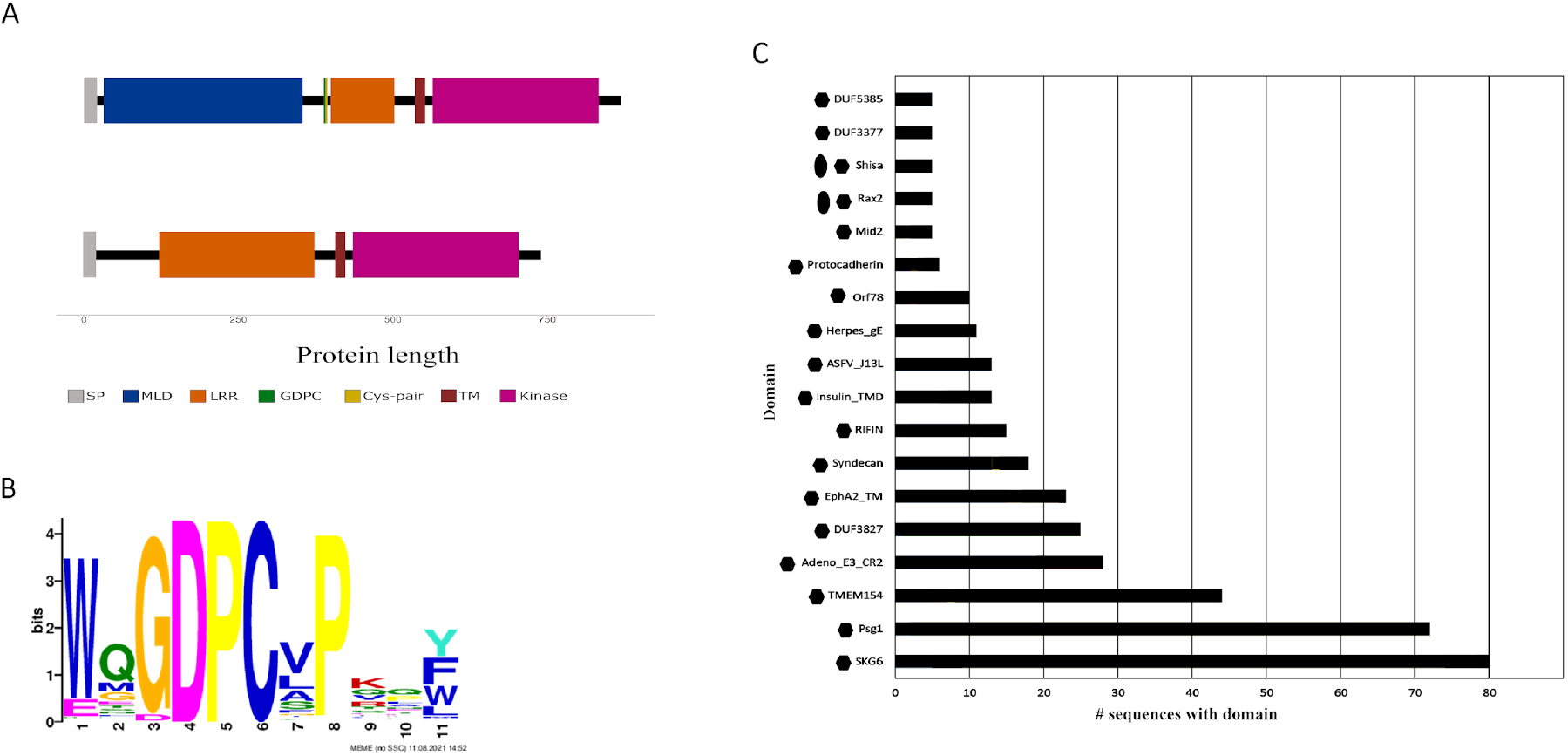
Domain architectures of LRR-I members across 112 species. General schematic of the two commonly found architectures (A) across 112 species. In (A) MLD = malectin-like domain, LRRs = leucine-rich repeats, TM = transmembrane domain, GDPC = a cleavage motif, Cys-pair = predicted disulfide bond, and SP = signal peptide. The two architectures are based on Arabidopsis FRK1 (top) and *P. patens* receptor Ppate_Pp3c16_23380V3.1.p (bottom). (B) Sequence logo of the WxGDCPxP motif found in the majority of LRR-I sequences. Uncommon domains (C) found within at least 5 sequences. Hexagons represent those found within the transmembrane domain, and ovals represent those found within the LRR domain.

We next investigated whether there were novel protein motifs found specifically within the ECDs of these proteins. In doing so, we found numerous highly conserved motifs within the extracellular region (Supplemental Figure 4). The most common non-LRR motif was the WxGDPCxP motif (GDPC, Figure 4B). Over 87% of all LRR-I receptors contain this motif located between the well-annotated malectin-like and LRR domains (Figure 4, Supplementary Figure 5-6). The majority of LRR-I sequences contain both the GDPC and MLD. Notably, however, the GDPC motif is mostly absent from receptors in LRR-I.1, LRR-I.6, and LRR-I.23, although LRR-I.6 and LRR-I.23 receptors do contain an MLD (Supplementary Figure 5).

## Discussion

Despite their well-documented roles in growth and immunity, the majority of plant LRR-RLKs still have no known biological function. Assigning functions to individual receptors is time-consuming and complicated by the large number of genes and potential for functional redundancy. We hope that the use of evolutionary and phylogenetic signatures to identify candidate LRR-RLKs that are likely to play roles in adaptation to stress may streamline this process. We therefore investigated the LRR-RLK repertoires of 112 plant and algal species to identify the subfamilies most likely to contain stress response receptors, which we hypothesize to be those with high levels of gene expansion particularly via tandem duplication (Qiao et al., 2019).

In accordance with previous reports (Fischer et al., 2016), our analysis showed varying rates of gene expansion and retention both between LRR-RLK gene subfamilies within a species, and in a single LRR-RLK subfamily between different species. While most subfamilies showed low levels of expansion across plant evolution and little variability between species, there were several subfamilies that showed increased gene number, variability in gene content across species, and potential asymmetric expansion. These features were especially prominent in subfamilies LRR-I, LRR-III, LRR-VIII-1, LRR-VIII-2, LRR-XI, and LRR-XII (Figure 1A).

These families display both high levels of expansion and between-species variability, and interestingly they all contain at least one member known to be involved in stress responses (Yeh et al., 2016; Le et al., 2014; Tang et al., 2015; Chinchilla et al., 2006; Aryal et al., 2023; Uemura et al., 2020).

We hypothesized that subfamilies that act broadly in stress responses would also show high levels of tandem duplication and subsequent gene retention. Duplication and retention provide genetic material for neofunctionalization and signal diversification to broaden the ability to detect biotic stress in particular (Hanada et al., 2008). While we observed evidence of tandem duplication in seven subfamilies in Arabidopsis, only three had more than one cluster of duplicated genes (Figure 1B). Subfamilies LRR-I, LRR-VIII-2, and LRR-XII are all greatly expanded via tandem duplication, with at least half of the total gene content found in duplicated clusters.

The only other subfamily to contain more than two tandem duplicated genes is LRR-VIII-1. It is interesting to note that LRR-I, LRR-VIII-1, and LRR-VIII-2 are the only three subfamilies that contain either a malectin or malectin-like domain and they have all been expanded via multiple tandem duplication clusters. Malectin domains are named after malectin, a conserved di-saccharide binding protein found in animal cells (Schallus et al., 2008, 2010). The ability to bind carbohydrates, combined with the expansion of the motif within plant genomes, led to speculation that receptor kinases containing these domains may act as cell-wall sensors (Yang et al., 2021). Subsequent work has shown that several of these receptors play roles in immunity signaling (HOK et al., 2011; Yeh et al., 2016; Chan et al., 2020; Yeh et al., 2015; Robatzek and Somssich, 2002). These findings would support the hypothesis that in Arabidopsis and other Brassicales, the expansion and retention of LRR-I genes is associated with their function in biotic stress response.

It is not clear why this expansion of LRR-I seems to come at the expense of expansion of the PRR-containing LRR-XII subfamily seen in most other lineages (Figure 2). Interestingly, the observed expansion in LRR-I coincides with several other genetic and life history changes in the Brassicales lineage. Many Brassicaceae are non-hosts to arbuscular mycorrhizae fungi (AM) or associate with only limited fungal species (Cosme et al., 2018). The Brassicaceae family also show a unique shoot-skewed expression pattern of the immune-related nucleotide-binding leucine-rich repeat receptors (NLRs). In contrast, other plant species preferentially express these resistance genes in the roots (Munch et al., 2018). The coincident loss of AM fungal association and NLR expression in the roots suggests that the Brassicaceae may have evolved a unique mechanism for protecting roots from pathogenic attack. Interestingly, most LRR-I genes in Arabidopsis display preferential expression in the root, positioning them perfectly to provide a non-specific immune response against a wide-variety of microbes (Supplemental Figure 7). In contrast, plants that depend upon associations with commensal fungi may require more specific root protection that can effectively identify pathogenic invaders while allowing beneficial associations. If true, this hypothesis would still raise the question of whether the loss of AM-fungal associations allowed the development of a non-specific root immunity, or whether the development of the immune response eliminated the possibility of commensal association.

To further investigate the possibility of LRR-I receptors gaining novel function, we also looked for the presence of other protein motifs or domains. The incorporation of such novel features can lead to functional diversity, a phenomenon recently shown for the intracellular NLR family of receptors (Van de Weyer et al., 2019). The majority of the newly identified domains span the transmembrane domain and have no known function. Given their localization, they may assist in receptor complex formation or signal propagation, a phenomenon that has been observed in other kinase systems (Arcas et al., 2020; Hubert et al., 2010; Li and Hristova, 2010).

In addition, we also identified the well-conserved GDPC motif located between the MLD and LRR regions in the majority of LRR-I sequences (Figure 4, Supplemental Figure 5, 6). This motif has been previously identified in LRR-RLK subfamilies LRR-I, LRR-IV, LRR-V, and LRR-VIII.1 (Dufayard et al., 2017), but we have identified it in a minority of LRR-VIII.2 receptors (210/1989 LRR-VIII.2 receptors, data not shown). This domain has been studied in depth in the case of the LRR-I member SYMBIOSIS RECEPTOR-LIKE KINASE (SYMRK). In *Lotus japonicus*, the GDPC motif acts as cleavage signal, resulting in the release of the MLD from the receptor (Antolín-Llovera et al., 2014). The release of the MLD allows the remaining receptor, containing only the LRR domains, to associate with a member of a family of Lysin-motif (LYS-M) containing receptors important for chitin recognition (Buendia et al., 2018). Given the conservation of the motif, we hypothesize that members of the LRR-I subfamily share a common regulatory mechanism dependent on cleavage and release of the MLD region.

While over 87% of all LRR-I receptors contain the GDPC motif, it is not found in LRR-I.1, LRR-I.6, or LRR-I.23 (Supplemental Figure 3). Although lacking the GDPC motif, LRR-I.6 and LRR-I.23 do contain the MLD (Supplemental Figure 3). In contrast, LRR-I.1 and LRR-I.2 are the only subclades lacking the MLD (Supplemental Figure 3). However, LRR-I.2 does contain the GDPC motif (Supplemental Figure 3). These sequences are found exclusively in Marchantia and raise the obvious question of what function cleavage at the GDPC motif may possibly serve in this context. The lack of the GDPC motif in LRR-I.1 but the presence in LRR-I.2 suggests the GDPC motif was acquired prior to the addition of the MLD domain to this subfamily of receptors. Lack of the GDPC motif in LRR-I.6 and LRR-I.23, then, may indicate that receptor cleavage is not always essential.

Through this work, we investigated the LRR-RLK family of receptor kinases across 112 species of plants and algae to find evidence of expansion. In our work, we found LRR-RLK subfamilies underwent asymmetric expansion across different lineages, with LRR-I being preferentially expanded in bryophytes and Brassicales. Furthermore, we found that the GDPC cleavage motif is found in the majority of LRR-I receptors, implying the motif is necessary for these receptors’ function. The work presented here provides new insights into the expansion and evolution of the LRR-I family. Further analysis of other lineages, including those within the gymnosperms, would give even more information on the evolution and expansion of the LRR-I family.

