## Supplementary material for "Differential expansion and retention patterns of LRR-RLK genes across plant evolution": Supp figures 1-7, table 1

**Supplemental Figures**

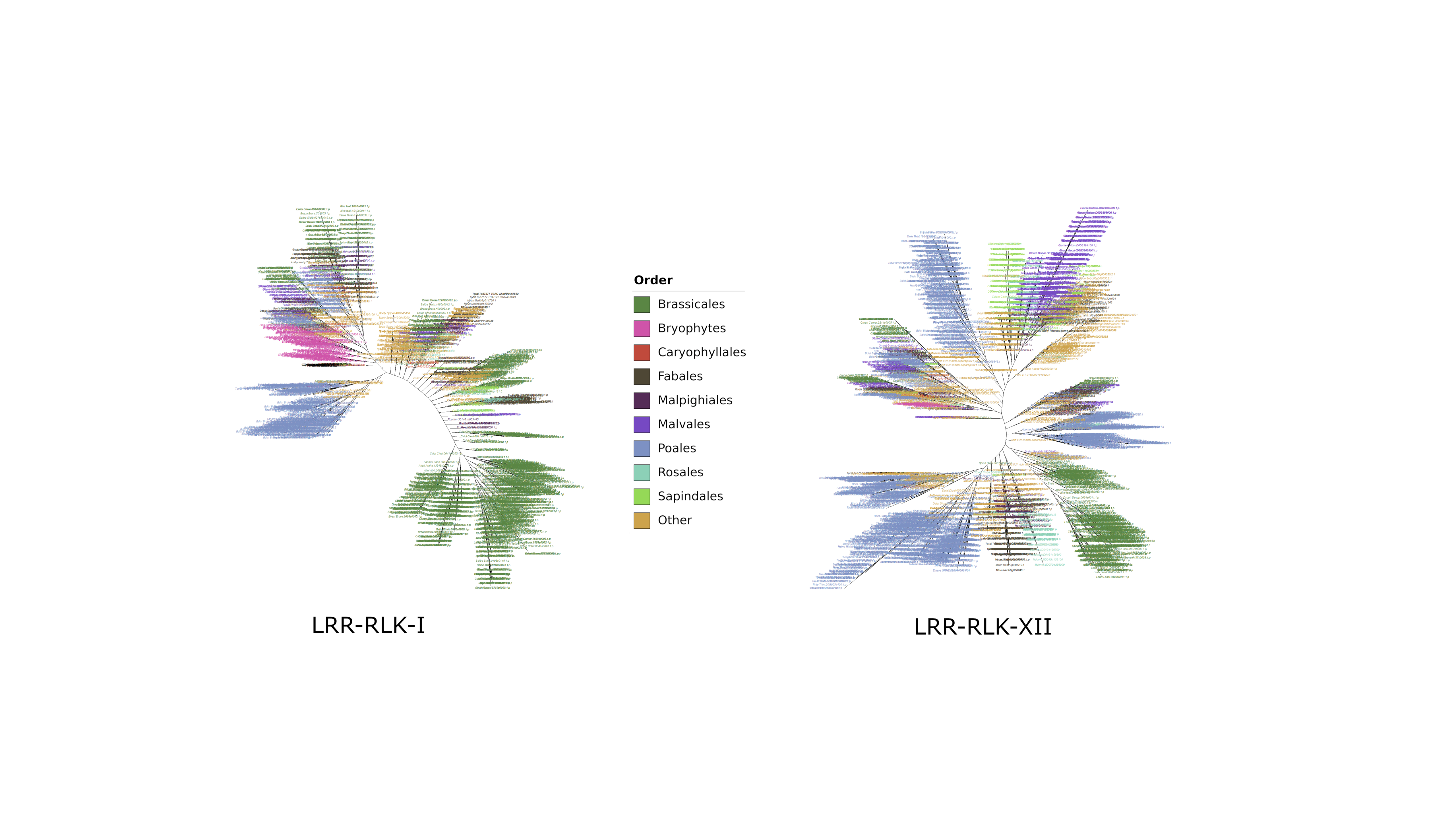

**Supplemental Figure 1. Unrooted distribution of LRR-I and LRR-XII genes across 112 species.** Trees were inferred using iqtree and model finder with default settings. Brassicales and Bryophytes have a higher incidence of LRR-I than LRR-XII, while the opposite is true for most other orders. Each leaf represents a protein. Proteins were labelled based on the plant order from which they are derived.

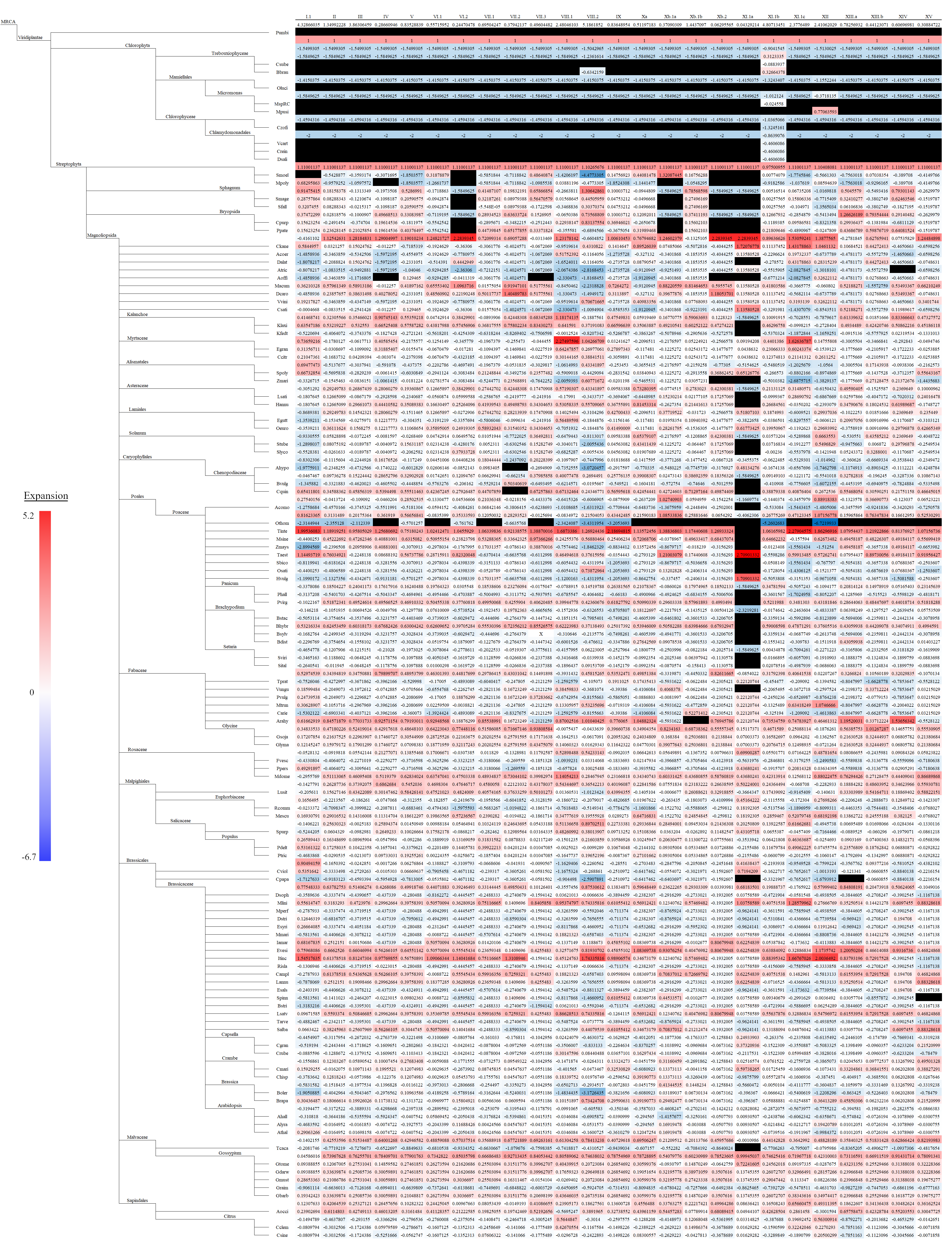

**Supplemental Figure 2. Expansion and contraction of the LRR-RLKs across 112 species compared to ancestral nodes.** The ancestral states were reconstructed using fastAnc for all LRR-RLK subfamilies. Values denoted here represent the rate increase or decrease compared to the most recent common ancestor at each node. Species are denoted in the tree based on in a hierarchical tree from NCBI taxonomy browser. Red values = increased expansion relative to the last common ancestor, blue = retraction relative to the last common ancestor, and white = no difference. Black values denote species with no known receptor in that subfamily.

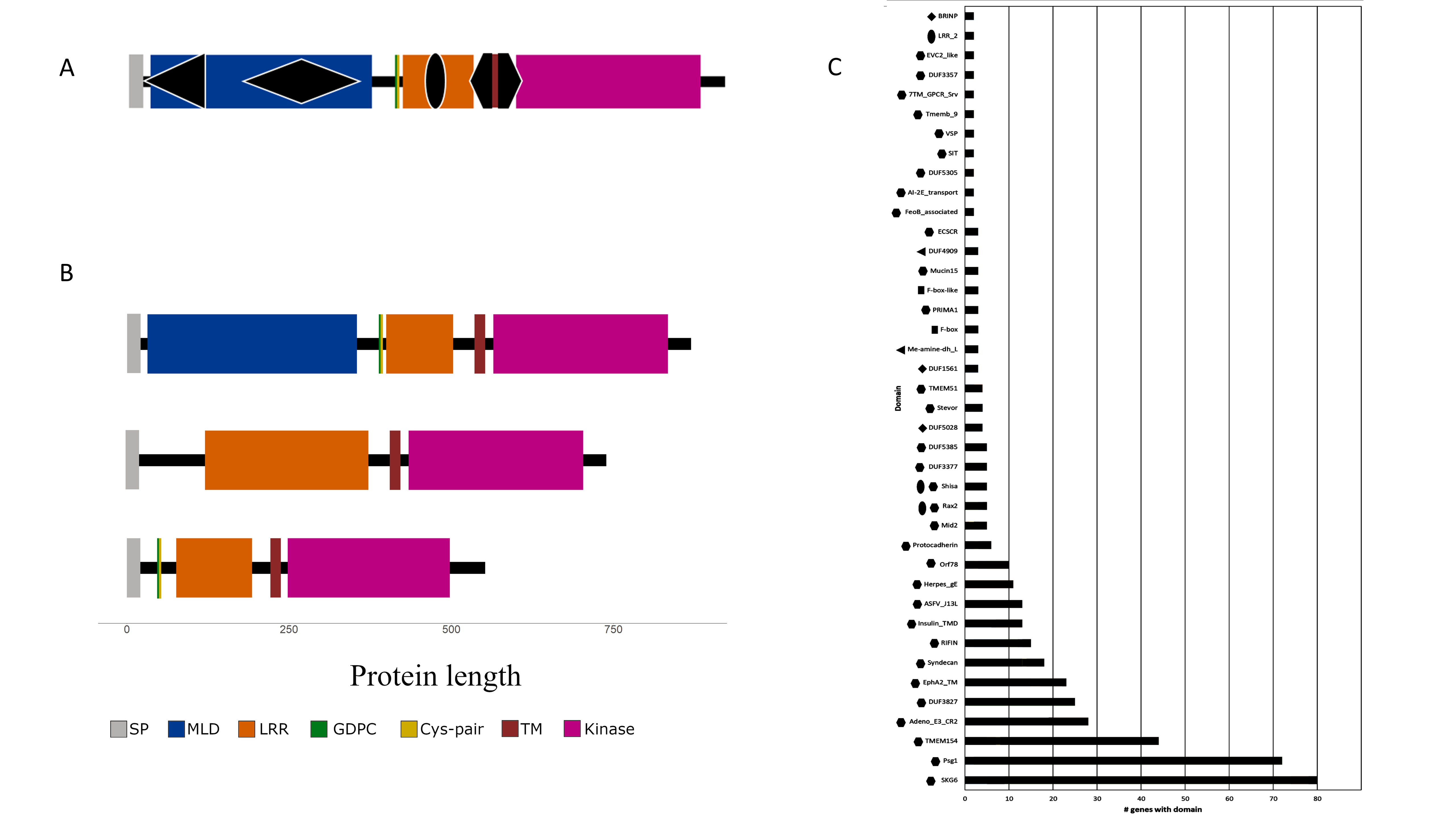

**Supplementary Figure 3. Analysis of uncommon domain architectures.** In (A) MLD = malectin-like domain, LRRs = leucine-rich repeats, TM = transmembrane domain, GDPC = a cleavage motif, Cys-pair = predicted disulfide bond, and SP = signal peptide. Domain architecture in (A) is represented by Arabidopsis FRK1. Alternate domain architectures, with the third, truncated architecture found in a smaller subset of proteins (B). Domains and motifs in (B) are mapped generally to representative receptors including Arabidopsis FRK1 (top), *S. magellanicum* receptor Smage_Sphmag04G082300.1.p (middle), and *B. distachyon* Bsylv_Brasy6G106300.1.p (bottom). Presence of GDPC motif and Cys-pairs in the truncated version (bottom) is variable across protein sequences. Domains presented in (C) represent those found in at least 2 genes across at least 2 different species. Symbols beside graph bars (C) represent general location the domain is found in (A).

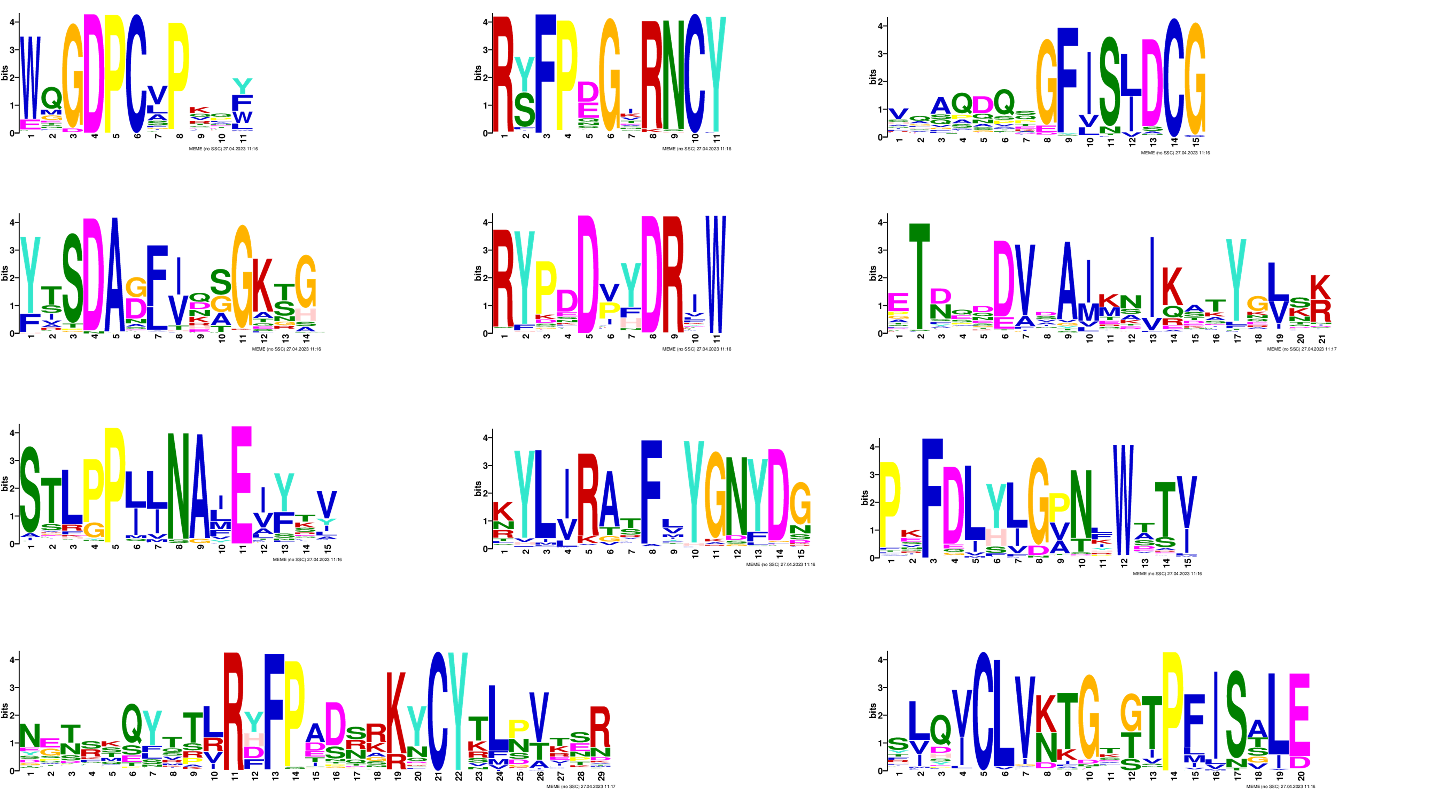

**Supplemental Figure 4**. Most significant non-LRR motifs found in LRR-I proteins. Bit score represents amino acid. Motifs present contain at least 5 amino acids with a bit score higher than 2. Motifs are collected from the top 100 identified sequence motifs using MEME suite on all LRR-I proteins. occurrence at each position.

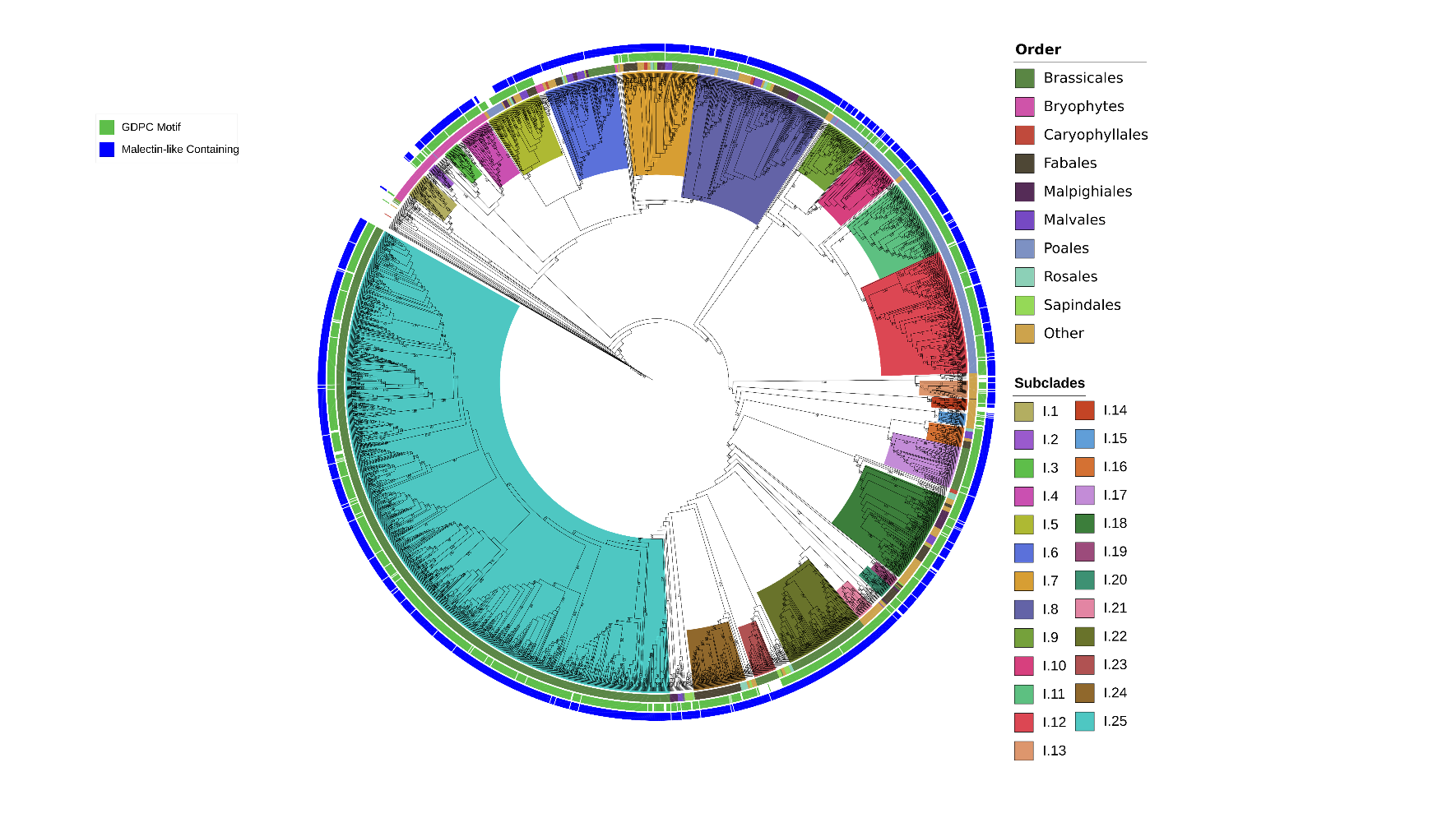

**Supplemental Figure 5**. **Motif analysis for GDPC and MLD domains**. Outer two rings (green and blue) have been added to the LRR-I rooted phylogeny from Figure 3. A green box in the middle ring denotes the presence of the GDPC motif, while a blue box in the outer ring denotes the presence of an MLD. Inner ring represents lineages of each sequence. Inner clades represent subclades.

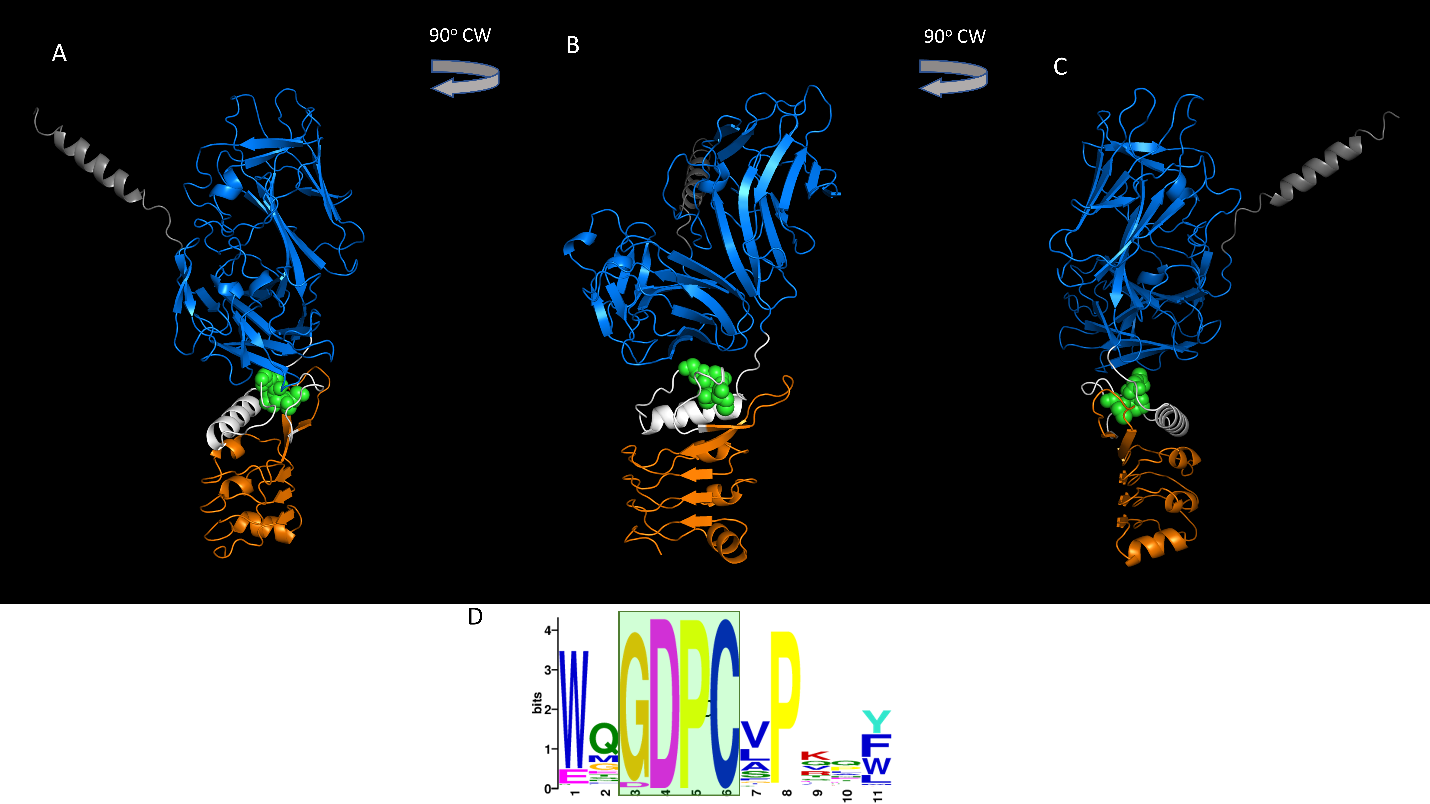

**Supplemental Figure 6**. **Schematic and labelled protein model of FRK1 (alphafold model AF-O64483-F1).** Grey = signal peptide, blue = malectin-like domain, orange = leucine-rich repeats, green spheres = GDPC motif, white = region with no known domain. Protein in (B) is rotated 90 degrees clockwise relative to (A), and (C) is rotated 90 degrees relative to (B). Sequence logo in (D) represents the GDPC motif.

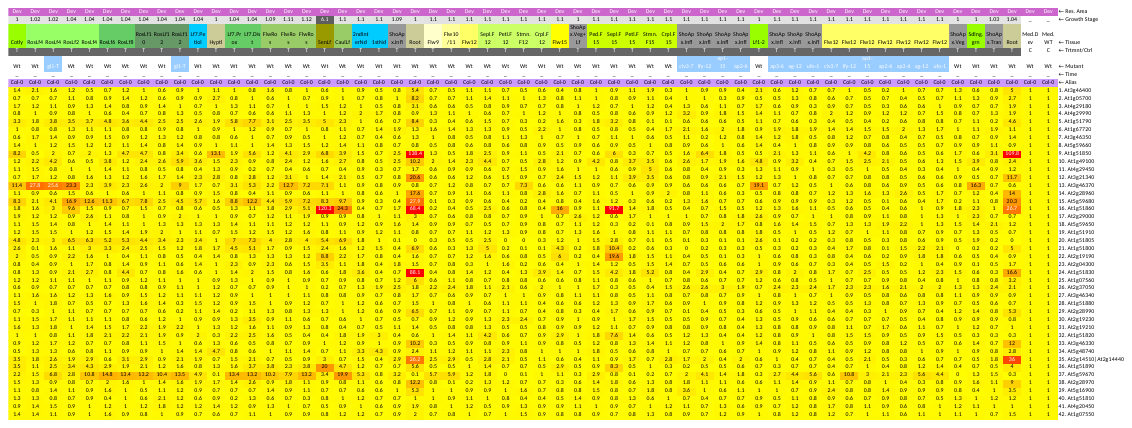

**Supplemental Figure 7. Expression analysis of 42 LRR-I genes from Arabidopsis across different developmental stages.** Red = highly expressed, yellow = lowly expressed relative to median values. Data and figure acquired from eNorthern at the BAR (Toufighi et al., 2005). Most genes show highest expression in roots.

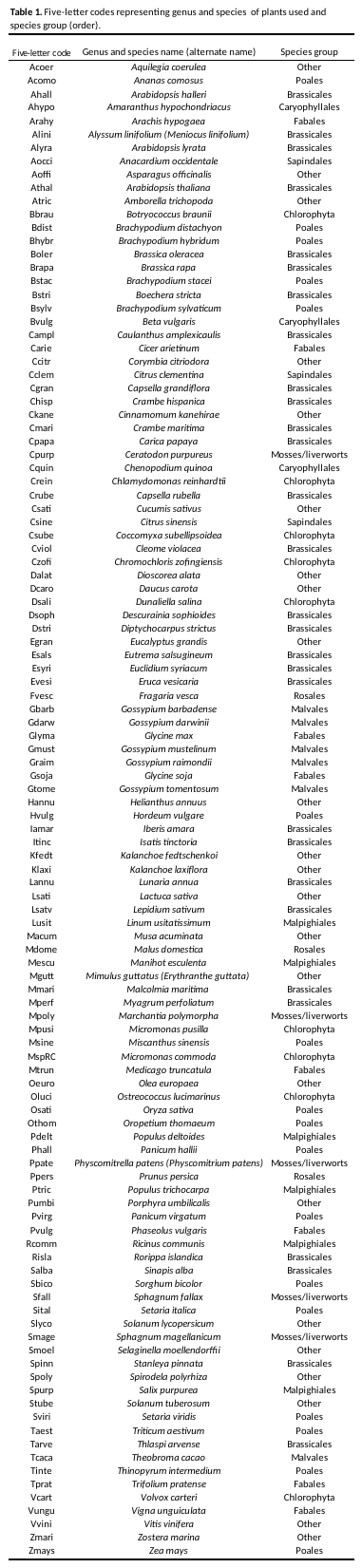
**Supplemental Tables**

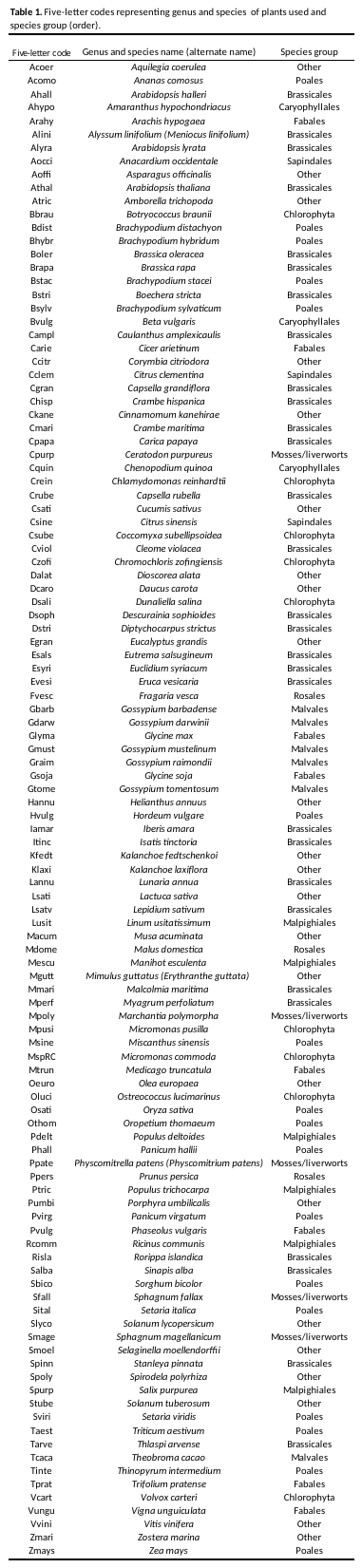

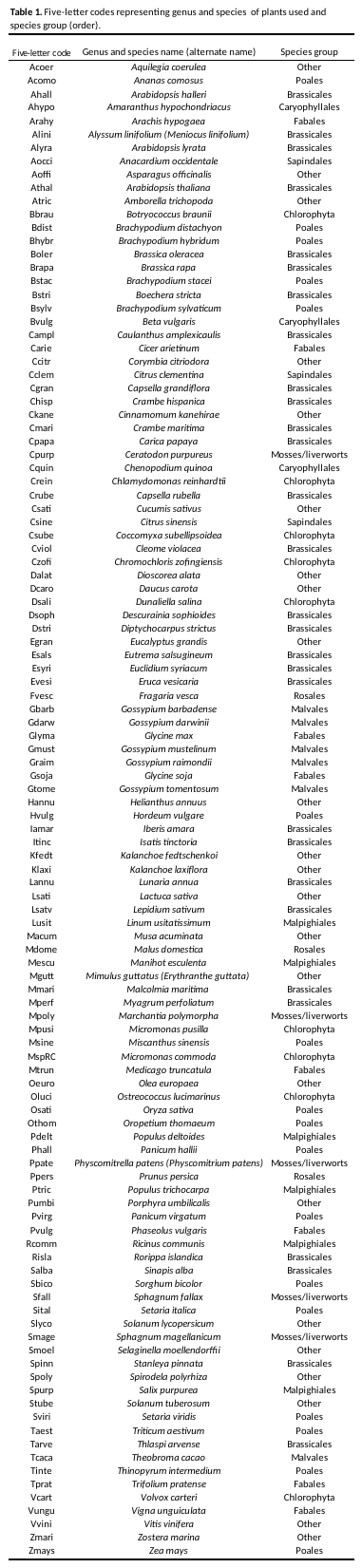
